# Examining the quality and fertilization competence of bovine ejaculate with low progressive motility – should we give it a chance?

**DOI:** 10.1101/2021.02.24.432701

**Authors:** Y. Li, D. Kalo, A. Komsky-Elbaz, Y. Zeron, Z. Roth

## Abstract

Spermatozoa progressive motility is positively correlated with fertilization competence. Bulls’ ejaculates with progressive motility lower than 50% are routinely rejected through the process of straw preparation, designated for artificial insemination of dairy cows. We examined the quality and fertility competence of ejaculates with relative low progressive motility (55–60%, n = 5; control) with those of very low progressive motility i.e. below the lower threshold, (20-45%, n = 5; rejected). Analysis revealed a lower volume for the control vs. rejected samples. Dip-Quick staining revealed a higher proportion of spermatozoa with abnormal morphology in the rejected group, in particular those with detached heads. Activation of spermatozoa with calcium ionophore, resulted by a lower proportion of activated spermatozoa in the rejected group. In addition, a higher proportion of spermatozoa with DNA damage were recorded in the rejected vs. the control samples. Following in-vitro fertilization, the proportion of oocytes that developed to the 2- and 4-cell stage embryos did not differ between groups. However, the proportion of embryos that further developed to blastocysts, was higher in the control group. Transcript abundance of selected genes in the blastocysts and the apoptotic index did not differ between groups, suggesting that the forming blastocysts were of the same quality. It is suggested that in specific cases, for example genetically superior bulls, ejaculates with very low progressive motility can be used for *in vitro* production of embryo. Further *in vivo* examinations, i.e. artificial insemination or transferring of embryos derived from these inferior ejaculates, might clarified this point.

## 1. Introduction

In modern dairy farms, artificial insemination (AI) is widely used for intensive reproductive management and it is the main approach used in genetic programs to improve herd merit (Meinert et al., 1992). As a single bull is generally used to breed numerous cows, each ejaculate should be of high quality.

In humans, male infertility due to asthenozoospermia is about 19% (Curi et al., n.d.). Assisted-reproduction treatments such as intrauterine insemination and *in vitro* fertilization (IVF), including intracytoplasmic spermatozoa injection, are used to overcome low motility and to improve couple infertility (Dyer et al., 2016). Nevertheless, these technologies are rarely used in livestock reproductive management. Other physiological key factors used to determine spermatozoa quality. Among these are spermatozoa motility and progressive motility (PM), which defined as the spermatozoa’s ability to move straight forward in a clearly defined direction (Li et al., 2016).

In bovine, PM has been shown to be associated with *in vivo* (Morrell et al., 2018) and *in vitro* (Kathiravan et al., 2011) fertilization competence. IVF with semen of high PM resulted in higher fertilization rate relative to that with low PM (>81% vs. <62%, respectively) (Li et al., 2016). Moreover, semen of very low PM (20-45%), is rejected through and not used for AI. This determinate approach may lead, in some cases, to a financial loss. For instance, ejaculate of a selected bull with desirable genetic traits can be rejected during the evaluation process, because of low physiological parameters, including PM.

The aim of the current study was to evaluated the quality of ejaculates with very low PM (<45%), and to associate physiological parameters and spermatozoa’s functions with *in vitro* embryonic development. Fresh ejaculates were assessed by computerized sperm-quality analyzer (SQA-Vb). Spermatozoa functions, including capacitation, acrosome reaction, binding to the zona pellucida and DNA fragmentation, were examined after cryopreservation. IVF was performed to evaluate fertilization competence and embryonic development. Finally, apoptotic index and the expression of genes involved in early embryonic development were examined to evaluate the quality of the developed blastocysts.

## 2. Materials and methods

### 2.1 Materials

All chemicals, unless otherwise specified, were from Sigma-Aldrich (Rehovot, Israel). The culture media i.e. oocyte maturation media (OMM), HEPES–Tyrode lactate (TL), sperm–TL, IVF–TL and potassium simplex optimized medium (KSOM) were prepared in our laboratory as previously described (Gendelman and Roth, 2012).

### 2.2 Semen collection

Ejaculates were collected during the winter at the Israeli Artificial Insemination Center (‘Sion’, Hafetz Haim, Israel) from 5-year-old Holstein Friesian working bulls. Semen was collected routinely twice a week. The first ejaculate on the collection day was taken for physiological characteristics by SQA-Vb (Orgal et al., 2012). Progressive motility served as the primary parameter to create two experimental groups. Ejaculates with low PM (55–60%; i.e., the lower cutoff for preparing straws) served as the control (n = 5; control) and were compared to ejaculates with PM lower than 20-45%, i.e., routinely rejected (n = 5; rejected). Ejaculates of representative bulls from the control and rejected groups were cryopreserved (Orgal et al., 2012) and evaluated for spermatozoa morphological, DNA integrity, acrosome reaction, hemizona assay (HZA) and IVF.

### 2.3 Spermatozoa purification

Frozen–thawed spermatozoa were purified by a Percoll gradient (45/90) followed by swim-up procedure (i.e., ‘Percoll’) as previously described (Parrish et al., 1995). Briefly, 3 mL of 45% Percoll (90% Percoll diluted 1:1 with Sperm-TL) was layered on 3 mL of 90% solution in a 15 mL tube. Spermatozoa (200 µL) were loaded onto the top of the gradient and then centrifuged for 10 min at 700*g*. Most of the non-motile spermatozoa were recovered at the interface of the Percoll layers, and the spermatozoa pellet at the tube bottom contained mostly motile spermatozoa. The pellet was immediately collected into a new 15 mL tube containing either 6 mL of NKM buffer (pH 7.4; 110 mM NaCl, 5 mM KCl, 20 mM MOPS [3-N-morphilino propanesulfonic acid, pH 7.4]) or SP-TALP medium (sperm-TL supplemented with 0.6% BSA, 1 mM sodium pyruvate and 0.2 mg/mL gentamicin) and allowed to swim up for 20 min at 38.5 °C. SP-TALP medium was used for acrosome reaction, hemizona assay (HZA) and IVF; a NKM buffer was used for assessment of spermatozoa viability and DNA integrity.

### 2.4 Assessment of spermatozoa functions

#### 2.4.1 Evaluation of spermatozoa morphology

None purified frozen–thawed samples were stained with Dip-Quick staining kit (Jorgensen Laboratories Inc., Loveland, CO) and spermatozoa were analyzed for morphological characteristics under an inverted microscope using NIS-Elements software (Nikon, Tokyo, Japan) at ×200 magnification. At least 200 spermatozoa were counted in each sample. The number of spermatozoa with detached head, bent or coiled tail or other abnormalities, such as small head, proximal or distal droplets or coiled midpiece, were recorded.

#### 2.4.2 Assessment of DNA integrity and spermatozoa viability

Frozen–thawed spermatozoa samples were purified and subjected to DNA integrity and viability assessment (Komsky-Elbaz et al., 2020; Komsky-Elbaz and Roth, 2018). DNA-compaction level was expressed as DNA fragmentation index (DFI), i.e. the ratio of the number of spermatozoa with fragmented DNA to the number of total spermatozoa. Spermatozoa viability was examined using the easyCyte kit no. 1 (Komsky-Elbaz and Roth, 2018). For both DFI and viability assessment a signal from 5000 events was read with a Guava easyCyte microcapillary flow cytometer, using CytoSoft software (Guava Technologies Inc., Hayward, CA, USA; distributed by IMV Technologies, L’Aigle, France).

#### 2.4.3 Assessment of acrosome reaction

Frozen–thawed spermatozoa were purified and subjected to acrosome reaction examination by a one-step staining method using fluorescein isothiocyanate-conjugated peanut agglutinin (FITC–PNA) (Orgal et al., 2012). For capacitation, purified spermatozoa (25 x 10^6^ spermatozoa/mL) were incubated with 0.5 mL SP-TALP for 4 h in a warm bath at 38.5 °C with a circulator for continuous mixing. At this point, 10 μL of sample was smeared on a slide to evaluate the proportion of spermatozoa with spontaneous acrosome reaction. In addition, acrosome reaction was induced by 2 μL of calcium ionophore A23187 dissolved in 5 mM DMSO, added to 1 ml of the sample. Samples were held at 38.5 °C for 20 min and 10 μL of ionophore-treated sample was smeared on a slide to evaluate the proportion of spermatozoa with reacted acrosome.

Slides were air-dried and dipped in absolute methanol for 10 min. Samples were then treated with FITC–PNA at a final concentration of 25 μg/mL and incubated for 35 min at room temperature in the dark. Acrosome reaction was examined under an inverted fluorescence microscope with FITC filter (excitation at 450–490 nm and emission at 515–565 nm) using NIS-Elements software. Spermatozoa without the acrosomal region or with bright green fluorescence in the equatorial zone were scored as acrosome-reacted. From each sample, at least 200 spermatozoa were analyzed randomly to evaluate the percentage of acrosome reaction.

#### 2.4.4 Hemizona assay (HZA)

Frozen–thawed spermatozoa were purified and subjected to HZA as previously described (Menzel et al., 2007). Bovine cumulus-oocyte complexes (COCs) were recovered from follicles and denuded by gentle vortexing in HEPES-TL supplemented with 0.3% (w/v) bovine serum albumin (BSA), 0.2 mM sodium pyruvate and 0.75 mg/mL gentamicin (HEPES-TALP) containing 1000 U/mL hyaluronidase, then placed in a droplet of IVF medium supplemented with 0.6% (w/v) essential fatty acid-free BSA, 0.2 mM sodium pyruvate, 0.05 mg/mL gentamicin and 0.01 mg/mL heparin (IVF-TALP) medium and equally microbisected by ultrasharp splitting blade (Bioniche Animal Health USA, Bogart, GA, USA). Each zona pellucida half was placed on a Petri dish, in a 100-µL spermatozoa-suspension droplet consisting of IVF-TALP medium with 5 × 10^5^ motile spermatozoa/mL, covered with mineral oil, and incubated for 4 h at 38.5 °C in 5% CO_2_. Thereafter, each zona pellucida half was removed and washed in HEPES-TALP medium. The number of spermatozoa tightly bound to the outer surface of the zona pellucida was counted using an inverted microscope (Nikon) at ×400 magnification.

The experiment was performed five times with a total of 25 zona pellucida halves per experimental group. In each run, thawed control and rejected spermatozoa were cultured with a matched zona pellucida pair, derived from the same oocyte.

### 2.5 In vitro production (IVP) of embryos

Cycles of IVF were performed as previously described (Kalo and Roth, 2011). Bovine ovaries were obtained from Holstein cows at a local abattoir. COCs were aspirated (Arav, 2001) and were incubated in OMM for 22 h. Matured COCs were co-incubated with frozen–thawed spermatozoa (control or rejected; ∼1 × 10^6^ cells) for 18 h. After fertilization, putative zygotes were denuded of cumulus cells and randomly placed in of 10 in 25-μL droplets of KSOM. All embryo droplets were overlaid with mineral oil and cultured for 8 d at 38.5 °C in an atmosphere of humidified air with 5% CO_2_ and 5% O_2_. Cleavage and blastocyst formation rates were examined 42 to 44 h and on days 7 and 8 after insemination, respectively. The proportion of hatched blastocysts was examined on day 9 post-fertilization and calculated out of the total number of 8-day blastocysts.

### 2.6 Evaluation of embryo quality

#### 2.6.1 Quantitative real-time PCR (qRT-PCR) assay

Samples of blastocysts were collected for qRT-PCR assay on day 7. Each sample consisted of three blastocysts, stored at −80 °C until RNA extraction. Poly(A) RNA was isolated using the Dynabeads mRNA DIRECT Kit according to the manufacturer’s instructions (Life Technologies, Carlsbad, CA, USA) and then subjected to reverse transcription (RT) using the Superscript II RT (Life Technologies) as previously described (Kalo and Roth, 2015).

Quantitative RT-PCR (qRT-PCR) was carried out with primers for *NANOG, OCT4, SOX2, PTGS2, DNMT1, TFAM* and *STAT3. YWHAZ* and *SDHA* were used as internal reference genes. The selected marker genes are important for proper embryonic development. The primers were derived from bovine sequences found in Genbank and specific primer pairs were designed using Primer 3.0 software.

#### NANOG

forward primer, 5’ACCAGCCTTGGAACAATCAG3’; reverse primer, 5’GTGGCCTCCAGATCACAGAC3’; 100bp.

#### OCT4

forward primer, 5’GTGAGAGGCAACCTGGAGAG; reverse primer, 5’ACACTCGGACCACGTCTTTC3’; 109 bp.

#### PTGS2

forward primer, 5’GAAATGATCTACCCGCCTCA3’; reverse primer, 5’TCTGGAACAACTGCTCATCG3’; 161 bp.

#### DNMT1

forward primer, 5’GCTTTACTGGAGCGATGAGG3’; reverse primer, 5’GAAGTCCTGGAGGCACTGAG3’; 103 bp.

#### STAT3

forward primer, 5’GTCGGCTACAGCCATCTTGT3’; reverse primer, 5’CCTGTCAACCCGTTTGTCTT3’; 118bp.

#### TFAM

forward primer, 5’CTGGTCAGTGCTTTGTCTGC3’; reverse primer, 5’CTAAAGGGATAGCGCAGTCG3’; 128 bp.

#### SOX2

forward primer, 5’GTCCTATGGTGCTGGATGCT3’; reverse primer, 5’GTTGATGTTCATGGCACAGG3’; 113 bp.

#### YWHAZ

forward primer, 5’GCATCCCACAGACTATTTCC3’; reverse primer, 5’GCAAAGACAATGACAGACCA3’; 124bp.

#### SDHA

forward primer, 5’GGGAGGACTTCAAGGAGAGG3’; reverse primer, 5’TCAACGTAGGAGAGCGTGTG3’; 112bp.

The qRT-PCR was conducted on a LightCycler^®^ 96 system (Roche, Basel, Switzerland) using a DyNAmo ColorFlash SYBR^®^ Green qPCR Kit (Finnzymes, Espoo, Finland) as previously described ^16^. Gene expression was quantified and analyzed by LightCycler 96 software ver. 1.1 and the ΔΔC_t_ method was used to calculate the relative expression of each gene (Kalo and Roth, 2015).

#### 2.6.2 Terminal deoxynucleotidyl transferase dUTP nick end labeling assay

The *in situ* cell death-detection kit from Roche (TUNEL; Indianapolis, IN) assay was used to detect DNA fragmentation in embryos as described previously (Kalo and Roth, 2011). In addition, samples were stained with 1 mg/mL Hoechst 33342 in phosphate-buffered saline with 1 mg/mL polyvinylpyrrolidone (PBS–PVP). The apoptotic cell ratio for each blastocyst was determined by calculating the number of TUNEL-positive blastomers out of the total number of blastocyst cells.

### 2.7 Statistical analysis

A dataset that consisting of one piece of data per bull was analyzed by JMP-13 (SAS Institute Inc., 2004, Cary, NC, USA) using one-way ANOVA followed by ‘Wilcoxon’ test, a non-parametric one-way procedure for data that are not normally distributed. Variables were: spermatozoa’s morphology, acrosome reaction, quantity of spermatozoa bound to the zona pellucida, and spermatozoa DNA fragmentation.

An overall comparison between experimental groups for embryonic development (cleavage rate, blastocyst-formation rate, hatching, and blastocyst apoptotic index) was performed by one-way ANOVA followed by Student’s t-test. Before analysis, data were arcsine-transformed.

Statistical analysis of qRT-PCR data was conducted on the expression of each gene within experimental groups relative to the control group (set to 1). Data are presented as means ± SEM. For all analyses, *P* < 0.05 was considered significant.

## 3. Results

### 3.1 Semen traits

Fresh ejaculates were examined by SQA-Vb and assigned to experimental groups (Table 1). The motility and progressive motility in the control (n = 5) and rejected (n = 5) ejaculates differed between groups (65.4 ± 0.6 vs. 46.4 ± 2.2 and 58.3 ± 0.5 vs. 37.8 ± 1.8 %, respectively; *P* < 0.014). The proportion of spermatozoa with normal morphology was higher in the control vs. rejected groups (*P* < 0.013; Table 1A). The velocity of progressively motile spermatozoa did not differ between the two groups but the volume of the control was lower than that of the rejected samples (*P* < 0.013; Table 1A). The motility, motile spermatozoa concentration (MSC) and progressive motile spermatozoa concentration (PMSC) were higher in the control than in the rejected samples (*P* < 0.053; Table 1B).

**Table 1.** Traits of fresh semen in control and rejected groups, evaluated by SQA-Vb machine. Presented are morphology, volume, concentration and velocity of progressively motile spermatozoa (**A**) and progressive motility (PM), motility, motile sperm concentration (MSC) and progressive motile spermatozoa concentration (PMSC) (**B**). Data are presented as mean ± SEM.

| **A** | Normal morphology | Volume | Concentration | Velocity of PM |
| --- | --- | --- | --- | --- |
| Group | (%) | (mL) | (10<sup>6</sup> cells/mL) | cells (μm/s) |
| Control | 80.0±1.2 | 5.0±0.4 | 1887.8±164.3 | 112.4±7.2 |
| Rejected | 69.8±1.4 | 9.4±0.6 | 1770.4±347.8 | 82.0±15.2 |
| *P*-value | 0.013 | 0.013 | 0.624 | 0.241 |

| **B** | PM | Motility | MSC | PMSC |
| --- | --- | --- | --- | --- |
| Group | (%) | (%) | (10<sup>6</sup> cells/mL) | (M/mL) |
| Control | 58.3±0.5 | 65.4±0.6 | 1235.9±108.4 | 1101.4±98.8 |
| Rejected | 37.8±1.8 | 46.4±2.2 | 797.4±151.3 | 688.7 ±144.6 |
| *P*-value | 0.014 | 0.014 | 0.053 | 0.053 |

### 3.2 Assessment of spermatozoa functions

#### 3.2.1 Spermatozoa morphology

Frozen samples were thawed and stained with Dip-Quick staining. The proportion of spermatozoa with abnormal morphology was higher in the rejected samples than in the control samples (*P* = 0.001; Table 2). In particular, the percentage of spermatozoa with detached heads (*P* = 0.001; Table 2, Fig. 1A).

**Table 2.** Frozen–thawed spermatozoa were stained with the Dip-Quick staining kit. Presented are total abnormalities, detached head, bent or coiled tail, and other abnormalities. Data are presented as mean ± SEM.

| Group | Total abnormal<br>(%) | Detached head<br>(%) | Bent or coiled tail<br>(%) | Other abnormal<br>features (%) |
| --- | --- | --- | --- | --- |
| Control | 19.1±1.0 | 5.5±3.1 | 12.4±3.6 | 1.1±0.5 |
| Rejected | 45.6±1.9 | 41.2±1.7 | 4.0±0.2 | 0.4±0.1 |
| *P*-value | 0.001 | 0.001 | 0.033 | 0.155 |

**Fig 1.**
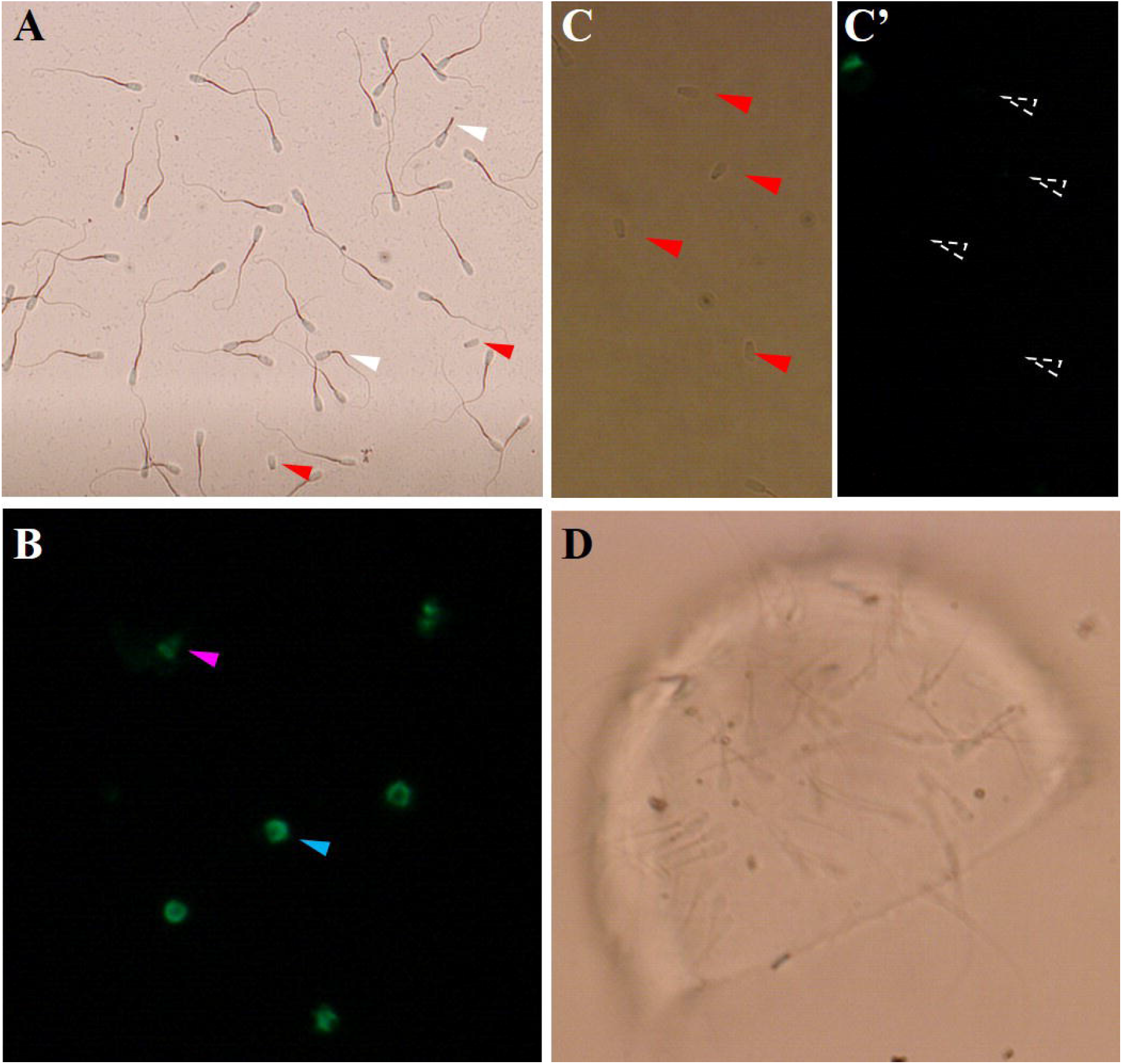
Representative images of spermatozoa from control and rejected samples (**A**) Fresh samples were purified and stained with Dip-Quick. White arrow, spermatozoa with bent or coiled tail; red arrow, spermatozoa with detached head. (**B**) Purified spermatozoa stained with FITC–PNA. Intact acrosome is characterized by bright staining over the acrosomal cap (blue arrow) and reacted acrosome is labeled in the equatorial segment (pink arrow). (**C**) Hemizona assay. Shown are spermatozoa bound to zona pellucida.

#### 3.2.2 Acrosome reaction

The percentage of spermatozoa with spontaneous acrosome reaction did not differ between the control and rejected samples (10.7 ± 2.6 vs. 7.4 ± 1.1%, respectively; *P* = 0.347). Following activation, the proportion of spermatozoa with reacted acrosome was lower in the rejected compared to control samples (46.1 ± 4.2 vs. 16.9 ± 2.1%, respectively; *P* = 0.009; Fig. 1B).

#### 3.2.3 Viability and DNA integrity

There was no difference in the percentage of viable spermatozoa between the control and rejected samples (55.5 ± 9.9 vs. 48.3 ± 6.4%, respectively). The proportion of spermatozoa with DNA damage was higher in the rejected samples relative to the the control, reflected by a higher DFI value (*P* < 0.0001; Fig. 2).

**Fig 2.**
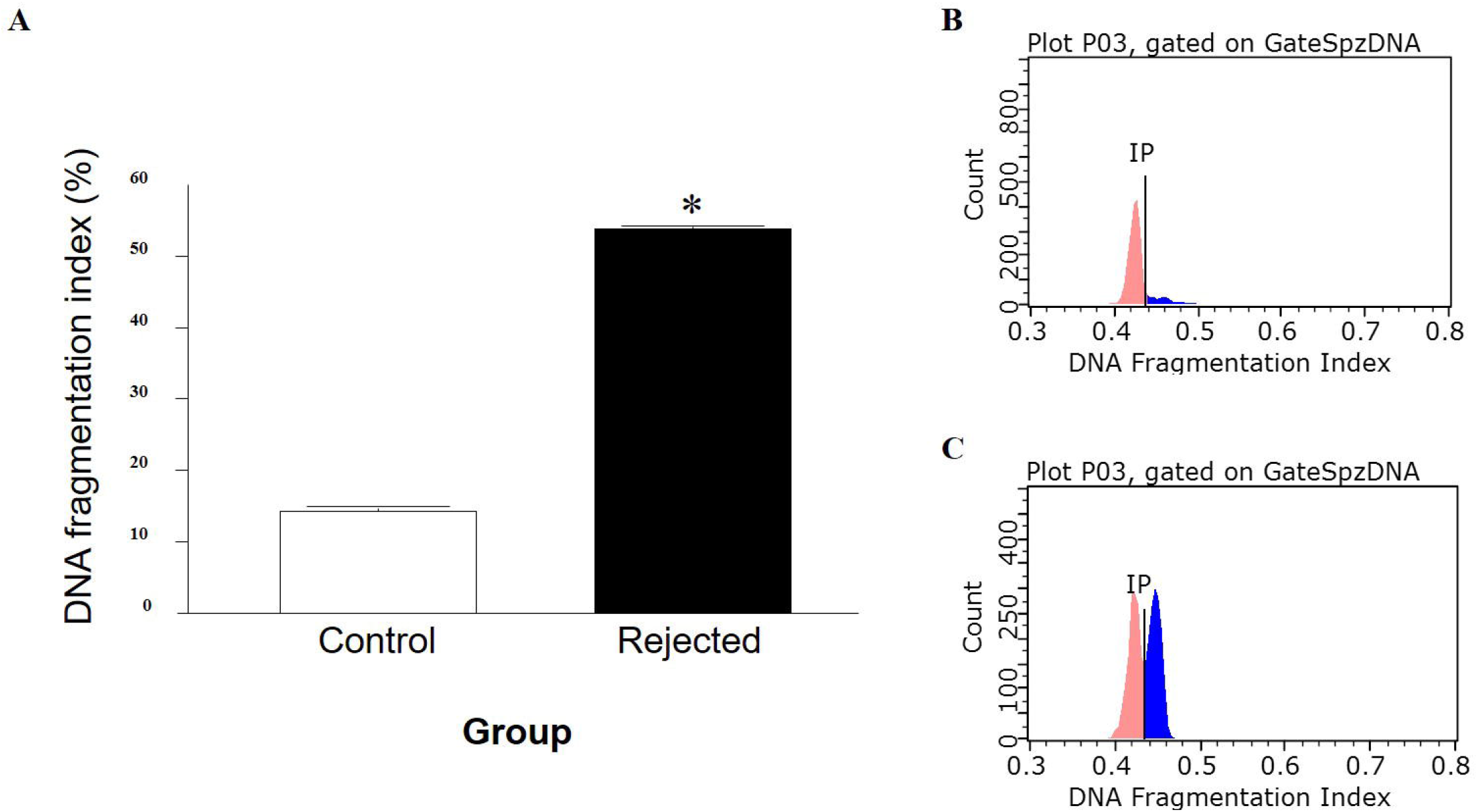
DNA integrity was evaluated in frozen–thawed samples purified by swim-up procedure. (**A**) Comparison of spermatozoa DNA-fragmentation index (DFI) in samples obtained from control and rejected groups. Acridine orange fluorescence cytograms for control group (**B**), with low proportion of spermatozoa with abnormal chromatin structure, and rejected group (**C**), with high proportion of spermatozoa with abnormal chromatin structure. Data are presented as the mean proportion of DFI ± SEM of the examined cells, calculated for four replicates, with 5000 spermatozoa/group examined.

#### 3.2.4 Hemizona assay

Difference in spermatozoon binding capacity was tested by HZA (Fig. 1C). The mean numbers of spermatozoa bound to the zona pellucida did not differ between control and rejected samples (143.4 ± 9.6 and 140.8 ± 12.4, respectively, *P* = 0.872).

#### 3.2.5 Cleavage and developmental competence

The experiment was performed six times with a total of 488 to 563 oocytes per group. The percentage of oocytes that were fertilized and cleaved to 2-to 4-cell embryos did not differ between the control and rejected groups (79.2 ± 2.2 and 77.2 ± 3.0 respectively, *P* > 0.05; Fig. 3A). However, the percentage of blastocyst stage embryos was higher for the control vs. rejected samples on both day 7 postfertilization (27.0 ± 2.3 and 19.9 ± 1.1, respectively, *P* < 0.05) and day 8 postfertilization (27.0 ± 2.3 and 19.9 ± 1.1, respectively, *P* < 0.05; Fig. 3B). Additional calculation of the proportion of blastocysts developed from cleaved embryos revealed similar differences for day 7 (34 and 26% for the control and rejected groups, respectively) and day 8 (28.2 and 18%, respectively).

**Fig 3.**
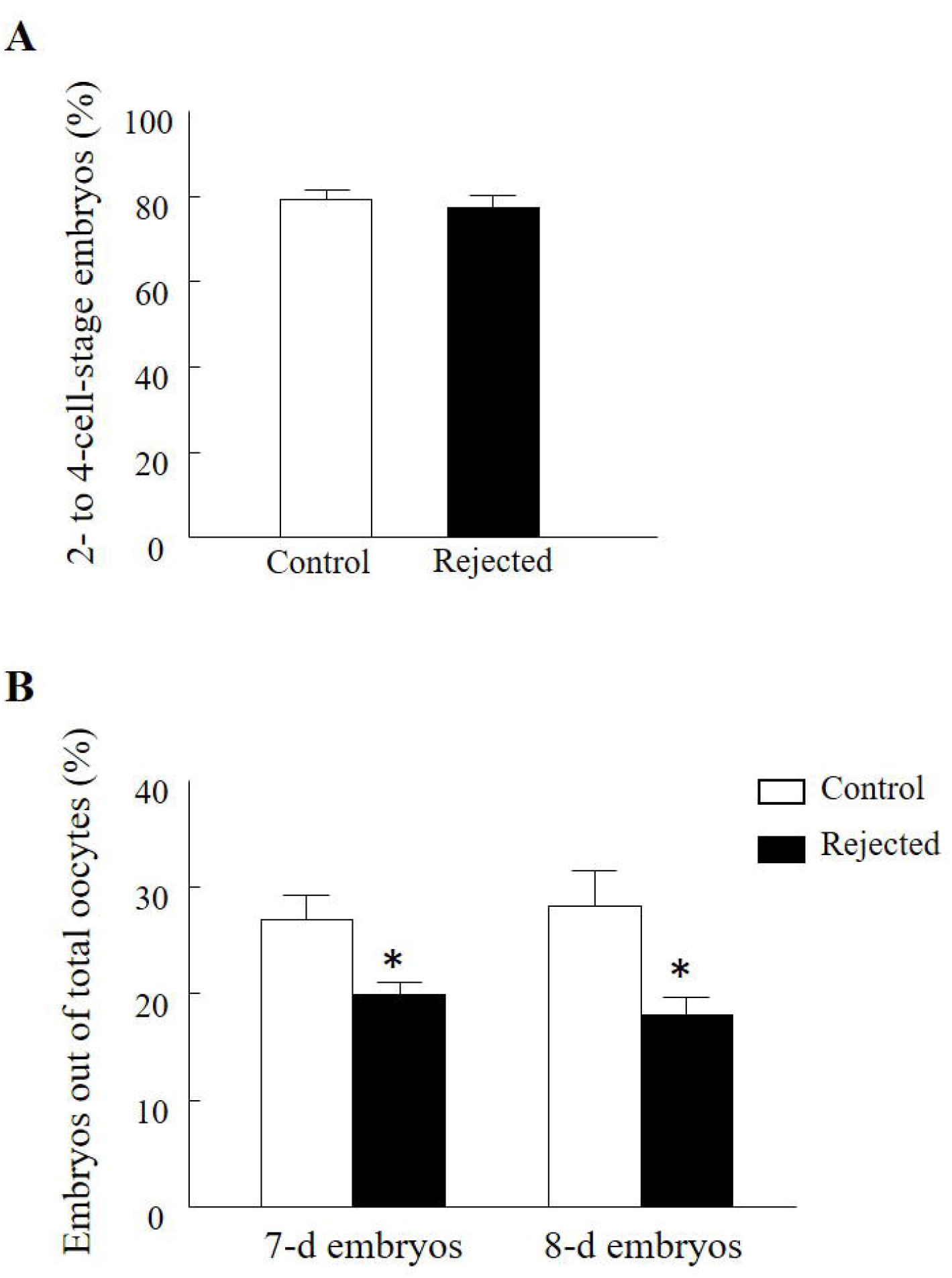
IVF competence of control and rejected samples. Cumulus-oocyte complexes were matured *in vitro*, then fertilized with either control or rejected purified spermatozoa and cultured for 7 d. Proportion of oocytes cleaved to 2-to 4-cell embryos, 42 to 44 h postfertilization (**A**), and proportion of embryos developed to the blastocyst stage on Days 7 and 8 postfertilization, calculated out of total oocytes (**B**). Data are presented as mean ± SEM. \**P* < 0.05.

#### 3.2.6 Embryo quality

The total cell number for control and rejected groups (62.3 ± 10.4 and 52.8 ± 13.6, respectively, *P* = 0.585; Fig. 4A) and the proportion of apoptotic cells (5.6 ± 0.7% and 8.6 ± 1.4%, respectively, *P* = 0.087; Fig. 4A, B) out of total cell number did not differ between the experimental groups.

**Fig 4.**
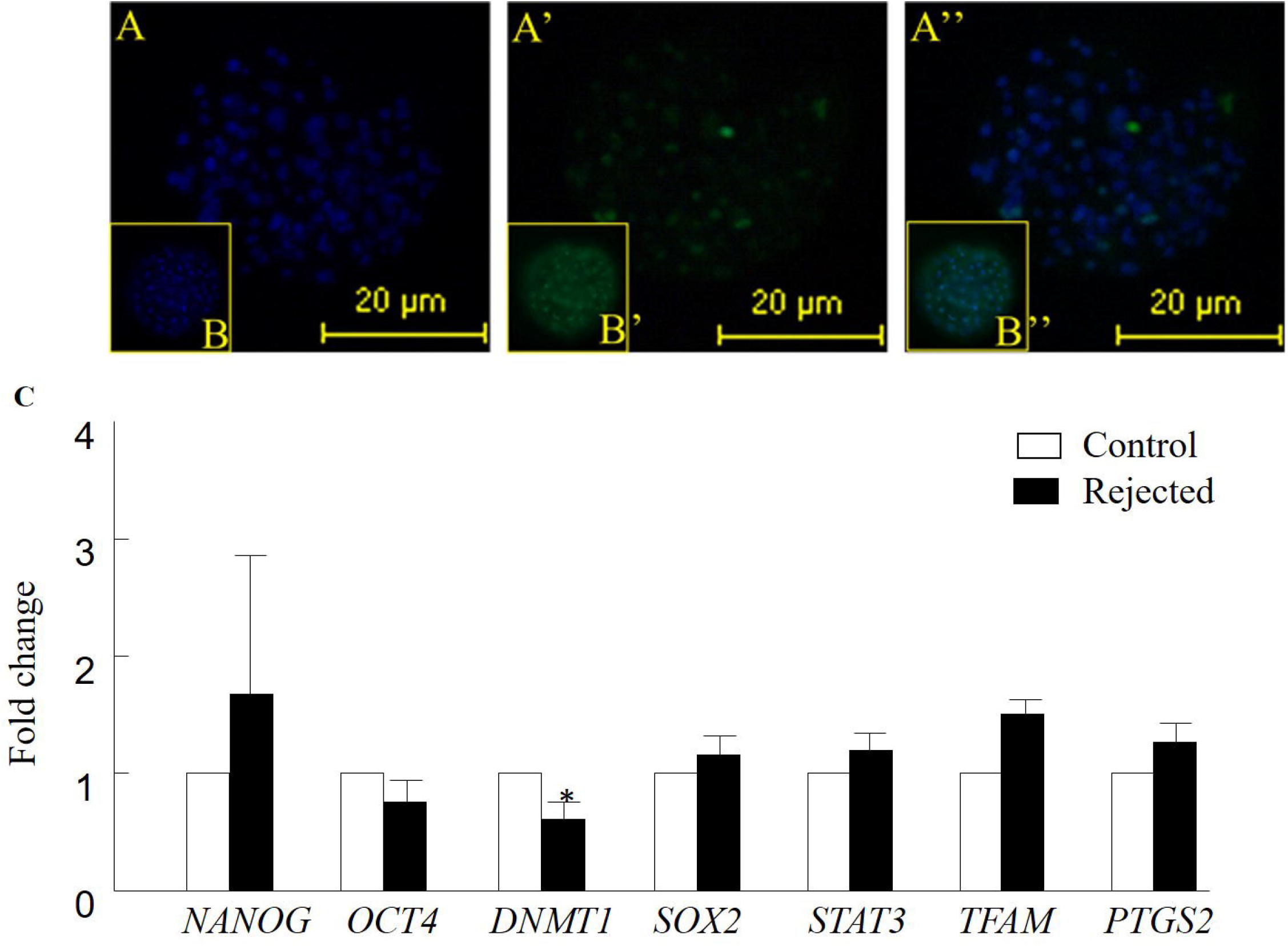
Quality examination of the developed blastocysts. Presented is 7-d blastocyst labeled with Hoechst 33342 staining (**A**, blue nuclei); TUNEL labeling (**A’**, bright green nuclei). Merged pictures of TUNEL labeling and Hoechst 33342 staining (**A’’**). **B, B’** and **B’’** are representative images of TUNEL-positive labeling. (**C**) qRT-PCR analysis of fundamental genes in blastocysts developed from control and rejected samples. Cumulus-oocyte complexes were matured *in vitro*, then fertilized with either control or rejected spermatozoa and cultured for 7 d. Transcript levels of *OCT4, SOX2, NANOG, TFAM, STAT3, DNMT1* and *PTGS2* were normalized to the geometric mean of *YWHAZ* and *SDHA* levels in blastocysts collected on day 7 postfertilization. qRT-PCR data were expressed relative to the control group. Data are presented as mean ± SEM. \**P* < 0.05.

The proportion of hatched blastocysts out of the total number of developed blastocysts did not differ between the control and rejected groups (3.6 ± 2.3% and 4.9 ± 2.5%, respectively, *P* = 0.714).

No alterations in the transcription abundance of *NANOG, OCT4, SOX2, PTGS2, TFAM* or *STAT3* in the developed blastocysts between experimental groups. However, a reduction in the expression level of *DNMT1* was recorded in blastocysts developed from the rejected group relative to the control (*P* < 0.05; Fig. 4C).

## 4. Discussion

A correlation between PM and IVF competence has been reported (Li et al., 2016). In particular, the proportion of embryos that fertilized, cleaved and developed to blastocysts was higher when fertilization was performed with ejaculates with high PM relative to those with low PM. Data from ‘Sion’ indicate that ∼7.8% of ejaculates are of low PM (<60%) and of these, ∼1.2% of the ejaculates are of very low PM (<45%); the latter are routinely rejected. Nevertheless, the fertilization potential of rejected ejaculates has never been examined in our system. Here we report that ejaculates of very low PM have fertilization capacity, expressed as a cleavage rate, similar to that of the controls. But, the proportion of the developed blastocysts was relatively lower than the control. With respect to the main question, “should we give these embryos a chance?”, it was found that the quality of embryos that developed from fertilization with semen of very low PM are of good quality at least according to their apoptotic index, transcript abundance of selected genes and the proportion of hatched blastocysts.

Ejaculate evaluation based on the PM parameter is routinely used at ‘Sion’, by using the SQA-Vb. Here we report an association between PM and sample volume; the latter was significantly higher in the rejected semen. While not clear, high ejaculate volume and reduced progressive motility have been documented for ejaculates collected during the summer (Majić Balić et al., 2012). In support, the percentage of ejaculates defined as being of inferior quality and therefore rejected, is higher in the summer vs. winter collection (Argov et al., 2007). In the current study, ejaculates were collected in the winter and therefore, therefore other stressors rather than thermal stress might underlie these alterations.

Spermatozoa motility has been reported to associated with embryonic developmental competence (Al Naib et al., 2011). It was reported that mild abnormalities in spermatozoa morphology have no influence on fertilization or cleavage rate, embryo quality, implantation, clinical pregnancy or live births (Shu et al., 2010). On the other hand, while moderate abnormalities in spermatozoa morphology did not affect fertility rate, it significantly decreased the proportion of good-quality embryos (Shu et al., 2010). Here we suggest that high proportion of spermatozoa with DNA fragmentation rather than morphological alterations might explain, at least in part, the reduced proportion of cleaved embryos that developed to the blastocyst stage. In the current study, high DFI was associated with reduced blastocyst formation. In support of this, the DFI calculated for spermatozoa with major-type abnormalities was significantly higher than that calculated for minor-type abnormalities (Shu et al., 2010). In addition, spermatozoa DNA fragmentation was reported to associate with aberrant embryo development (Tesarik et al., 2004). Although, DNA-repair mechanisms within the oocyte are activated upon fertilization and can repair DNA damage (Fernández-Díez et al., 2016), the oocyte has limited capacity to repair high levels of DNA fragmentation after fertilization and during early embryonic development (Wright et al., 2014).

In general, ejaculates collected from Holstein–Friesian bulls, the percentage of total abnormal morphologies and head defects did not exceed 20% and 8%, respectively (Hoflack et al., 2007). In the current study, about 19.1% of the spermatozoa in the control samples expressed abnormal morphology and only 5.5% were defined as having detached heads, indicating that despite their relatively low progressive motility (60%), the control samples were in the normal range. On the other hand, about 45.6% of the spermatozoa in the rejected ejaculates showed abnormal morphology. Of these, 41.2% were defined as having detached heads. A high proportion of spermatozoa with detached heads has been previously reported, without any association to genetic predisposition, imperfect spermiogenesis or pathogens (Siqueira et al., 2010).

Spermatozoa morphology is an important parameter in predicting fertility competence however, the association with embryonic development is not yet clear. In bovine, decreased embryonic development has been associated with morphological abnormalities in the spermatozoa, mainly with changes in head morphology (Walters et al., 2005). In humans, pregnancy rate per cycle decreases significantly as the normal morphology decreases, in association with increased abortion rate per pregnancy (Jinno et al., 1992). In the current study, spermatozoa morphology did not affect zona pellucida-binding capability. In particular, the excessive proportion of spermatozoa with detached heads in the rejected samples did not affect the proportion of spermatozoa bound to the zona pellucida. In addition, neither the proportion of spermatozoa with spontaneous acrosome reaction nor that of those that were artificially activated, differed between the groups. Taken together, it is reasonable to conclude that ejaculates of very low PM (i.e. the rejected group) expressed fair fertilization activity in vitro. In support of this, the fertilization ability of these ejaculates, expressed as cleavage rate into 2- and 4-cell stages, was relatively high (80%) and did not differ between groups.

Nevertheless, the proportion of cleaved embryos that further developed to blastocysts was lower when IVF was performed with rejected samples. In humans, embryos with poor morphology have been associated with a low proportion of morphologically normal spermatozoa (Parinaud et al., 1993). However, this was not the case in the current study. Neither the morphology of the blastocysts nor the proportion of hatched embryos differed between the experimental groups. Moreover, the total cell number and the proportion of TUNEL-positive cells did not differ between control and rejected groups, suggesting that embryos were of the same quality. In addition, the expression of selected genes, excluding that of *DNMT1*, did not differ between groups. DNMT1 is involved in the DNA-methylation mechanism by maintaining the DNA-methylation pattern during replication (Moore et al., 2013). In addition, DNMT1 is important for early embryonic development since its depletion in mice induces embryonic lethality between embryonic d 8.0 and 10.5 (Li et al., 1992). Nonetheless, further molecular and genetic examination should be performed to confirm these findings.

## 5. Conclusions

In summary, rejected ejaculates were found to express impaired morphological characteristics. In addition, a high proportion of DNA fragmentation was noted in the rejected semen. While these alterations did not affect fertilization ability, the proportion of developing blastocysts was lower for the rejected-than the control semen. Nevertheless, the developed blastocysts seem to be of good quality. It is suggested therefore, that in specific cases, for example genetically superior bulls, ejaculates of very low quality can be used for AI or producing embryos *in vitro* for embryo transfer. This suggestion warrants further examination *in vivo*.

## Author contributions

**Ying Li**: Data curation, Formal analysis, Methodology, Writing - original draft. **Dorit Kalo**: Methodology, Writing-review & editing. **Alisa Komsky-Elbaz**: Methodology, Writing-review & editing. **Yoel Zeron**: Resources. **Zvi Roth**: Conceptualization, Data curation, Supervision, Project administration, Writing – review & editing.

